# The differing roles of flavins and quinones in extracellular electron transfer in *Lactiplantibacillus plantarum*

**DOI:** 10.1101/2022.07.29.502109

**Authors:** Joe G. Tolar, Siliang Li, Caroline M. Ajo-Franklin

## Abstract

*Lactiplantibacillus plantarum* is a lactic acid bacteria that is commonly found in the human gut and fermented food products. Despite its overwhelmingly fermentative metabolism, this microbe can perform extracellular electron transfer (EET) when provided with an exogenous quinone, 1,4-dihydroxy-2-naphthoic acid (DHNA) and riboflavin. However, the separate roles of DHNA and riboflavin in EET in *L. plantarum* has remained unclear. Here we seek to understand the role of quinones and flavins for EET by monitoring iron and anode reduction in the presence and absence of these small molecules. We found that either addition of DHNA or riboflavin can support robust iron reduction, indicating electron transfer to extracellular iron occurs through both flavin-dependent and DHNA-dependent routes. Using genetic mutants of *L. plantarum*, we found that flavin-dependent iron reduction requires Ndh2 and EetA, while DHNA-dependent iron reduction largely relies on Ndh2 and PplA. In contrast to iron reduction, DHNA-containing media supported more robust anode reduction than riboflavin-containing media, suggesting electron transfer to an anode proceeds most efficiently through the DHNA-dependent pathway. Furthermore, we found that flavin-dependent anode reduction requires EetA, Ndh2, and PplA, while DHNA-dependent anode reduction requires Ndh2 and PplA. Taken together, we identify multiple EET routes utilized by *L. plantarum* and show that the EET route depends on access to environmental biomolecules and on the extracellular electron acceptor. This work expands our molecular-level understanding of EET in Gram-positive microbes and provides additional opportunities to manipulate EET for biotechnology.

**Importance:** Lactic acid bacteria are named because of their nearly exclusive fermentative metabolism. Thus, the recent observation of EET activity - typically associated with anaerobic respiration - in this class of organisms has forced researchers to rethink the rules governing microbial metabolic strategies. Our identification of multiple routes for EET in *L. plantarum* that depend on two separate redox active small molecules expands our understanding of how microbes metabolically adapt to different environments to gain an energetic edge and how these processes can be manipulated for biotechnological uses. Understanding the role of EET in lactic acid bacteria is of great importance due to the significance of lactic acid bacteria in agriculture, bioremediation, food production, and gut health. Furthermore, the maintenance of multiple EET routes speak to the importance of this process to function in a variety of environmental conditions.

## Introduction

Microorganisms inhabit a rich diversity of environmental niches including those with highly limited resources. Consequently, some bacteria have developed unique metabolic strategies to survive in resource poor environments. One such strategy is extracellular electron transfer (EET) that allows cells to achieve redox balance by transferring electrons out of the cell to terminal electron acceptors in their environment via an electron transfer network.^1,2^ Historically, this ability was thought to be limited to Gram-negative microbes because of the presence of a large, insulated cell wall in Gram-positive microbes.^3^ However, it was recently discovered that *Listeria monocytogenes*, a Gram-positive opportunistic human pathogen, utilizes a flavin-based EET pathway^4–6^ that is linked to a genetic locus (FLEET locus). This allowed us and others to identify other Gram-positive microbes capable of EET, including *Enterococcus faecalis* and *Lactiplantibacillus plantarum*.^5,7–9^ At present, however, it is not understood how exclusively fermentative microbes, such as *L. plantarum* and *E. faecalis*, can utilize and maintain an EET pathway, especially when a traditional respiratory system is not maintained. Elucidating this phenomenon and its influence on organismal physiology is of great importance due to the prevalence of these Gram-positive bacteria in agriculture, bioremediation, food production, gut health, and opportunistic human infections.^10–13^

Prior work in other bacteria provides insight into the possible role of different redox active small molecules, such as quinones and flavins, in EET. In *L. monocytogenes*, electrons travel from intracellular NADH to a membrane-confined quinone pool via Ndh2, a type II NADH dehydrogenase, then to an extracellular flavolipoprotein, PplA, and finally to terminal electron acceptors including ferric iron and electrodes.^5^ A number of studies have also found quinone derivatives can support EET by functioning as extracellular electron shuttles between microbes and insoluble electron acceptors.^14,14^ Additionally, flavins are known to mediate or support extracellular electron transfer in *Clostridium, Shewanella, Geobacter* and other microbes. ^16-21^.

We recently showed that the genus *Lactiplantibacillus* has a highly conserved FLEET locus and many of its members exhibit EET activity.^9^ The FLEET locus encodes a type-II NADH dehydrogenase (Ndh2), flavin transport proteins (FmnA/B and ATPase1/2), membrane demethylmenaquinone (DMK) synthesis proteins (DmkA and EetB/DmkB), and electron transfer proteins (PplA and EetA).^7–9^ An exogenous menaquinone (MK) precursor, 1,4-dihydroxy-2-naphthoate (DHNA), was added to allow the microbe to synthesize MK or DMK. When both an electron acceptor (ferric iron or an artificial electrode) and DHNA were provided, EET activity was observed in *L. plantarum*. Closer analysis of the FLEET locus across *L. plantarum* strains found that transposon insertions in either *ndh2* or *pplA* caused a loss of EET activity.^9^ While establishing the importance of riboflavin and DHNA in EET in *L. plantarum*, the precise role of these molecules in extracellular electron transfer remains unclear.

Through a series of electrochemical assays coupled with genetic knockouts, this study identifies and characterizes two distinct pathways for EET in *L. plantarum*. We found that both riboflavin and DHNA can independently support robust EET, but differ in the extent to which they support iron reduction and anode reduction. Here we show functional Ndh2, PplA, and EetA are required for flavin-dependent EET, while only Ndh2 and PplA are necessary for DHNA-dependent EET. Additionally, we found that DHNA can act as a robust electron shuttle between microbes and a carbon felt electrode, while riboflavin is less efficient as an electron shuttle under the same conditions. Overall, we have identified distinct electron transfer mechanisms that utilize commonly found redox-active small molecules and that may have evolved to provide interchangeable energetic means to adapt to different environments

## Results

### L. plantarum uses either DHNA or riboflavin to support extracellular electron transfer

Prior work has demonstrated *L. plantarum* requires PplA to reduce iron, but not for electrode reduction. The differential requirement of PplA could be caused by the difference in electron acceptor or because cells were prepared and grown differently depending on the assay used.^9^ Recently, studies of extracellular electron transfer in other organisms that contain the FLEET locus, *L. monocytogenes*^5^ and *E. faecalis*^8,22^, suggest such requirements may be dependent on growth conditions. Thus, we sought to understand how growth conditions affect whether DHNA and riboflavin are necessary to support iron reduction.

To test this, we first grew cells in mMRS alone or in mMRS supplemented with ferric ammonium citrate and DHNA. Cells from each pre-assay condition were washed, incubated in media with 20 µg/mL DHNA or 2 µg/mL riboflavin, then assayed for their ability to reduce iron(III) oxide nanoparticles under anaerobic conditions (SI Figure 1). We found that pre-assay media composition had no effect on iron reduction in wells lacking DHNA or riboflavin as well as those containing DHNA (Figure 1A). Conversely, we found that pre-assay media supplemented with DHNA+iron supported robust iron reduction activity in riboflavin-containing assay conditions (Figure 1A). We also found that supplementing the pre-assay media with only DHNA was sufficient to enable the riboflavin-dependent iron reduction (SI Figure 2). These results show that when grown with ferric ammonium citrate and DHNA, *L. plantarum* can use either riboflavin or DHNA to reduce iron. Since supplementing pre-assay media with both ferric ammonium citrate and DHNA resulted in the most robust iron reduction, all of the following experiments were conducted under those conditions.

**Figure 1:**
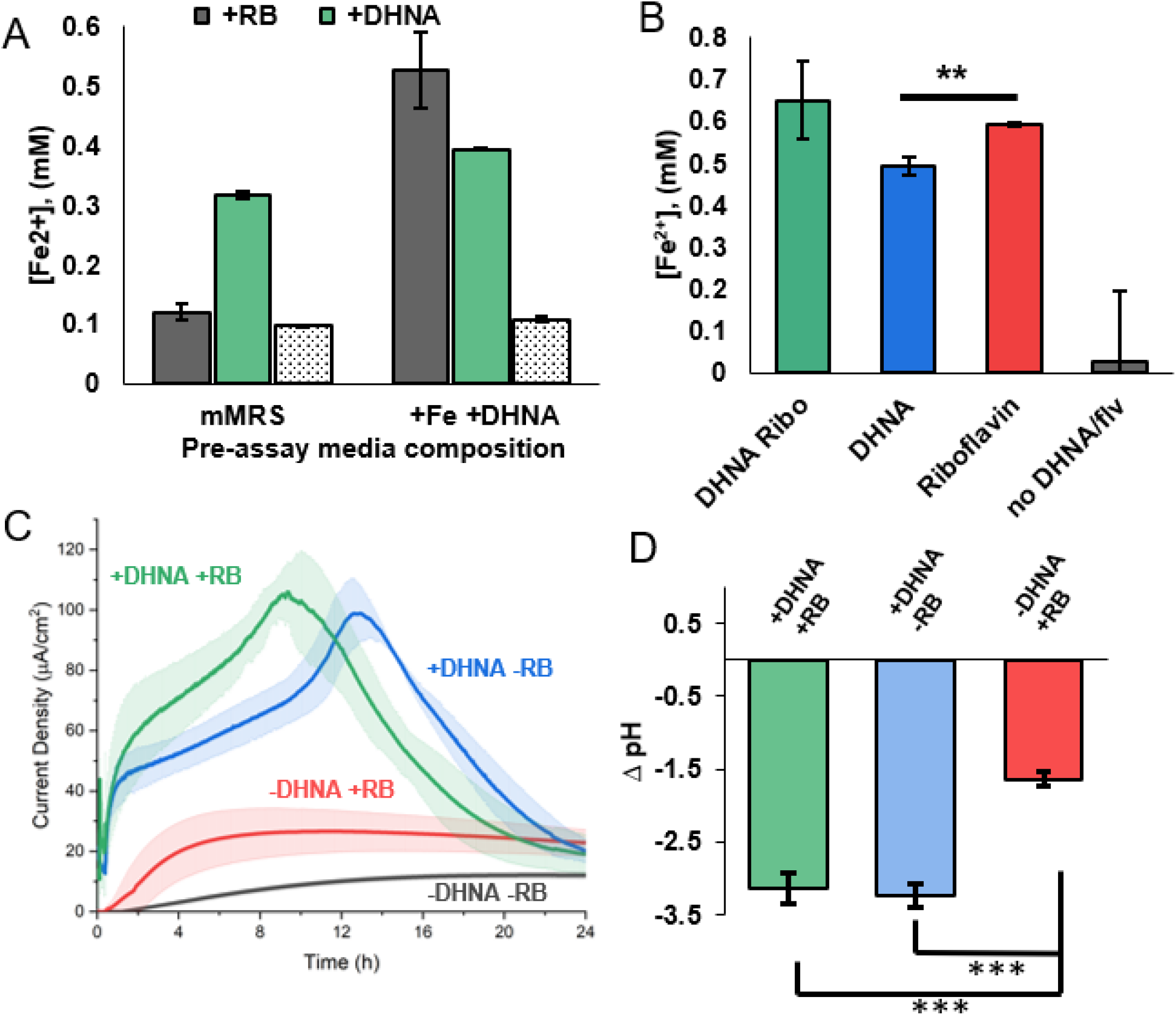
DHNA or flavins can support EET activity in distinct ways. (A) 500 Concentration of Fe^2+^ produced from ferric oxide nanoparticles after 24 hours anaerobic incubation in PBS + 20 μg/mL Mannitol (grey) alone or supplemented501 with DHNA (green), Riboflavin (patterned). Pre-assay media composition indicates overnight culture source contained exogenous DHNA and ferric ammonium citrate or was unsupplemented. (B) Concentration of Fe^2+^ produced from ferric oxide after 24 hours anaerobic incubation in PBS + 20 μg/mL Mannitol with DHNA and Riboflavin (green) compared to DHNA (blue) or riboflavin (red) alone. (C) *L. plantarum* was grown in microaerobic 3-chamber bioelectrochemical reactors for 24 hours on an electrode poised at 0.2V vs Ag/AgCl (3M KCl). (D) After 24h, the media pH was measured and is reported as the change in pH compared to the starting pH of 7.4. Chronoamperometric measurements were taken every 36s. Error bars show SD. Statistical significance calculated with a t-test (***= p<0.001).

**Figure 2:**
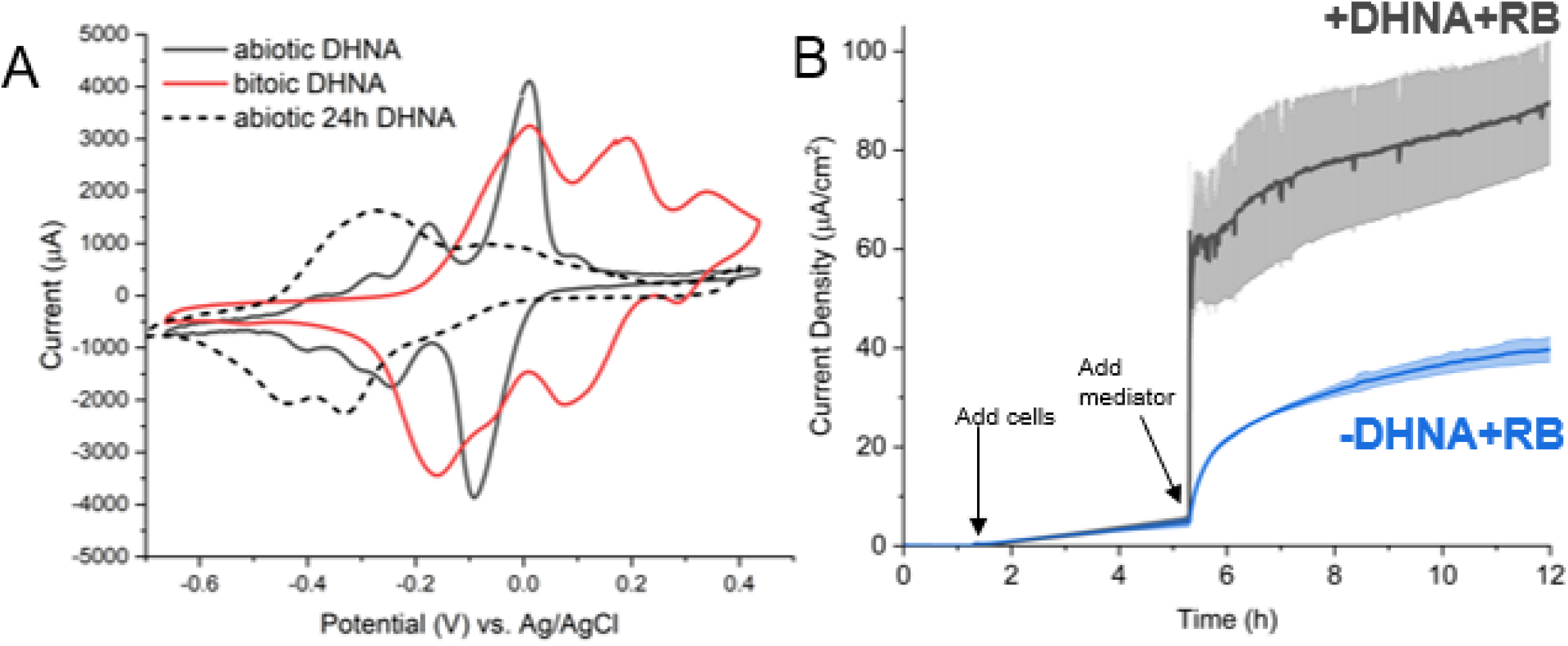
DHNA exhibits redox shuttle characteristics. (A) Cyclic voltammetry was performed prior to the addition of cells and 24h after the addition of cells. (B) *L. plantarum* was grown in microaerobic 3-chamber bioelectrochemical reactors for 3 hours on an electrode poised at 0.2V vs Ag/AgCl (3M KCl) before Riboflavin (blue) or DHNA and riboflavin (gray) was injected. Current density measurements were taken every 36s for 12h. Error bars show SD.

Because DHNA and riboflavin both have midpoint potentials more negative than iron (III) oxide, it is possible that their effects on iron reduction could be additive. To determine whether iron reduction by riboflavin and DHNA could be additive, we measured iron reduction by *L. plantarum* in media with DHNA and riboflavin, DHNA alone, riboflavin alone, and media lacking both. As expected, *L. plantarum* lacking both riboflavin and DHNA did not reduce iron (Figure 1B). In contrast, combining riboflavin and DHNA did not have an additive effect on iron reduction (Figure 1B). Together, these results indicate that iron reduction by *L. plantarum* proceeds at approximately equal rates through a DHNA- or flavin-dependent pathway.

While iron reduction provides researchers with a method for large scale screening, EET activity can be measured with exquisite temporal resolution via the reduction of a poised electrode. Many studies have shown that EET activity can vary between iron reduction and anode reduction.^9,22^ Because extracellular iron reduction is supported by riboflavin alone, we decided to test if riboflavin can support anode reduction when reactor conditions mimic iron reduction conditions. To do this, we replaced iron (III) oxide with an anode biased at +0.20 V vs Ag/AgCl. We found that riboflavin can support anode reduction, but the maximum current density is ∼4 fold lower than media containing DHNA or DHNA and riboflavin (Figure 1C). In DHNA supplemented media, the current rapidly increased to reach a maximum current density within 10-13 h and then gradually decreased. In riboflavin-supplemented media, the current gradually increased and reached a sustained plateau by ∼4 h. These data indicate that riboflavin and DHNA can both support EET to iron or an anode. However, the differing rates and time-evolution of flavin-dependent and DHNA-dependent EET to a poised electrode suggest this EET may occur by different mechanisms.

While it is clear that DHNA supports more robust current production than riboflavin, the physiological implications remain unclear. EET activity in *L. plantarum* was recently found to accelerate fermentative metabolism and resulted in a drop of extracellular pH.^9^ To see if the DHNA- and flavin-dependent routes equally contribute to the fermentative flux, we measured the pH of the media after 24 h and found that the change in pH was significantly smaller in the riboflavin containing media compared to media containing DHNA (Figure 1D). These data suggest that DHNA-dependent EET is more closely linked to the fermentative flux observed by Tejedor-Sanz et al^9^ than flavin-dependent EET.

### DHNA acts as a robust, reversible electron shuttle

Having established that DHNA supports both anode and iron reduction by *L. plantarum*, we next sought to characterize the mechanism of DHNA-dependent anode reduction. Our first step was to distinguish between two ways DHNA could support EET : either by acting intracellularly as a precursor for DMK, which is an membrane-confined electron carrier in EET in *L. monocytogenes*^5^ or, DHNA could be acting as a mediator, much like its close analog 2-amino-3-carboxy-1,4-naphthoquinone (ACNQ).^23^ To evaluate the two hypotheses, we performed cyclic voltammetry in reactors containing freshly dissolved DHNA, DHNA that had been in the reactor for 24 h, and DHNA incubated with cells for 24 h. We found that freshly dissolved DHNA has two clear redox peaks (−175 mV and -11mV) that correspond to literature values (Figure 2A). Interestingly, under abiotic conditions over 24 h, DHNA exhibited a redox shift to ∼ -320 mV, which is in line with its conversion to ACNQ (Figure 2A)^23^. Most profoundly, we found that DHNA in the presence of bacteria exhibits the -11 mV peak and an additional peak at ∼ +196 mV (Figure 2A). These data strongly strongly suggest that DHNA is acting as a mediator.

We next performed a pulse-chase experiment to observe the real-time bioelectrochemical response to the addition of DHNA. After a 3-4 h acclimation period for injected cells, either riboflavin or DHNA+riboflavin were added and the current production was sampled for 12 h. The addition of DHNA+riboflavin resulted in a large, immediate jump in current production, whereas the addition of just riboflavin resulted in a much slower increase in current production (Figure 2B). To quantify the time required to utilize DHNA, we calculated the amount of time required to reach half the maximal current, t_1/2max_, for two concentrations of DHNA and found that DHNA induced a response with a half-time of ∼14s at 20 μg/mL and ∼800s at 0.2 μg/mL (Table 1). This time scale is most consistent with a freely diffusible molecule^24^, providing additional support for DHNA’s role as a mediator.

**Table 1.** Quantitative characterization of EET reaction rate to DHAN and Riboflavin. *L. plantarum* was grown in microaerobic 3-chamber bioelectrochemical reactors for 3 hours on an electrode poised at 0.2V vs Ag/AgCl (3M KCl) before the indicated concentration of riboflavin or DHNA was injected. Current density measurements were taken every 36s for 24h. Half time was determined as ½ the time at which maximum current was achieved. Error bars show SD.

| *Condition* | *Max current density*<br>*($\mu\text{A}/\text{cm}^2$)* | *Half-Time to max*<br>*(seconds)* |
| --- | --- | --- |
| *0.2 $\mu\text{g}/\text{mL}$ DHNA* | *15.86 ±2.82* | *792* |
| *20 $\mu\text{g}/\text{mL}$ DHNA* | *70.36 ±1.55* | *14.4* |
| *0.2 $\mu\text{g}/\text{mL}$ Riboflavin* | *24.32 ±0.22* | *3096* |
| *2 $\mu\text{g}/\text{mL}$ Riboflavin* | *45.02 ±1.80* | *2988* |
| *20 $\mu\text{g}/\text{mL}$ Riboflavin* | *55.17 ±5.26* | *684* |

As an additional test, we performed a media swap experiment where DHNA was supplied to cells for ∼16h and the cells were collected, washed and inoculated into fresh media lacking DHNA. As before, DHNA enabled current production, but swapping the media to remove DHNA abolished it; the addition of fresh DHNA restored current production (SI Figure 3). Taken together, these data indicate DHNA can act as a reversible redox mediator that receives electrons and can donate them to a poised electrode.

### DHNA-dependent EET requires Ndh2 and PplA

With the identification of a DHNA-dependent route for anode reduction that appears to utilize DHNA as an extracellular electron shuttle, we wanted to characterize which FLEET genes were required to support anode reduction. In a prior study, under different experimental conditions, Ndh2 and PplA were shown to be necessary for iron reduction, but only Ndh2 was required for anode reduction.^9^ To identify which FLEET genes were required to perform DHNA-dependent EET under these conditions, knockouts of Ndh2, PplA, EetA/B, DmkA, and DmkB were created using the CRISPR-Cas9/RecET system.^25^ When these mutants were tested for their ability to produce current, Δ*dmkA*, Δ*dmkB*, Δ*eetA/B* showed similar current output compared to WT (Figure 3A). Conversely, Δ*ndh2* and Δ*pplA* had a ∼2-fold decrease in current compared to WT (Figure 3A). These data confirm the importance of Ndh2 under multiple experimental conditions. Additionally, these data show PplA can be required for anode reduction under non-growth conditions, whereas it was not required under conditions that promoted cell growth. Given the importance of *ndh2*, we also deleted *ndh1*, a second, type-II NADH dehydrogenase in *L. plantarum*. This deletion had no effect on current production compared to WT (Figure 3B), indicating *ndh1* is not involved in EET. Thus, the DHNA-dependent anode reduction proceeds primarily through Ndh2 and PplA with DHNA acting as a redox mediator (Figure 6).

**Figure 3.**
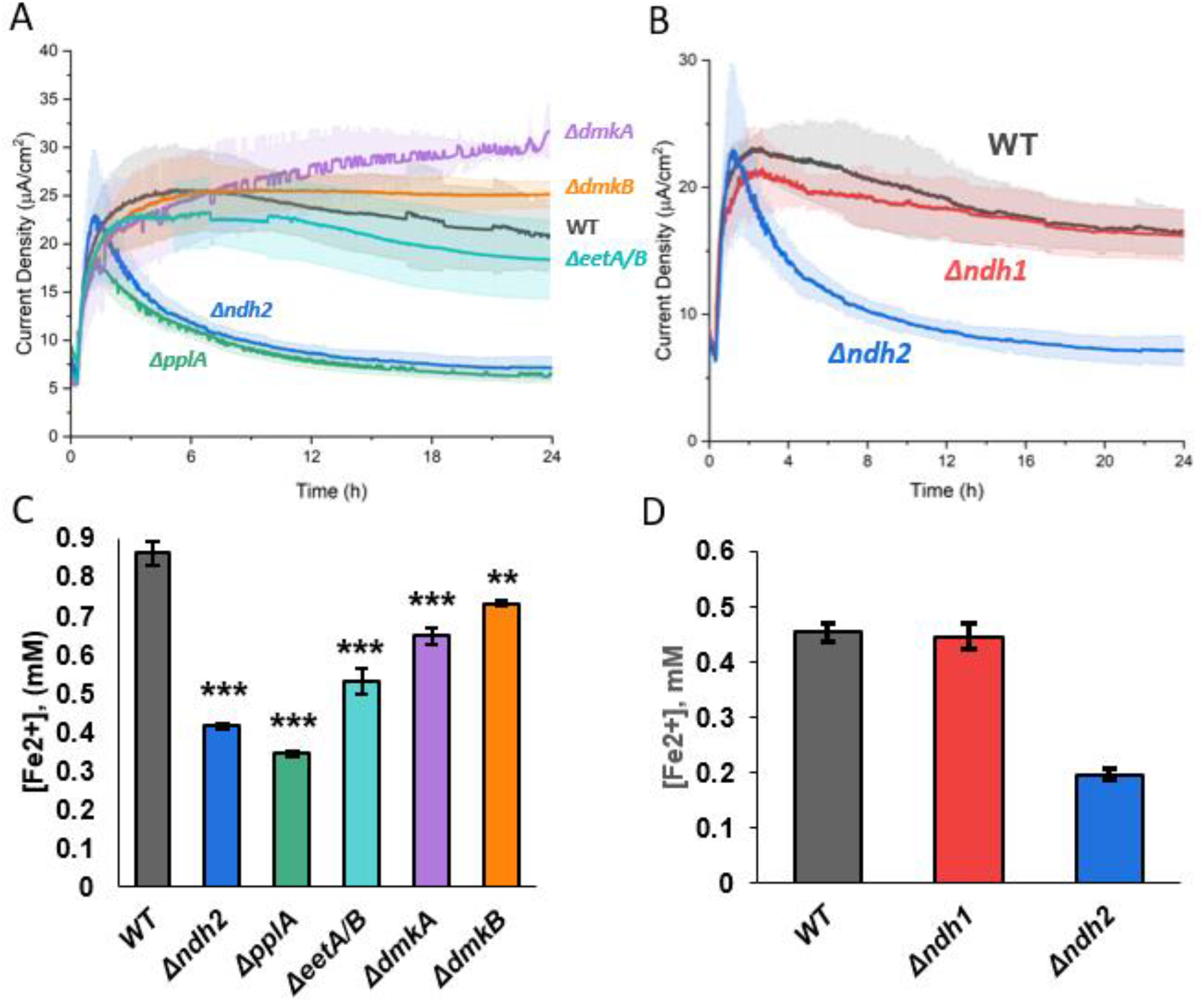
DHNA-dependent EET utilizes Ndh2 and PplA to reduced extracellular electron acceptors. Chronoamperometry was performed in 2-chambered 3 electrode bioreactors. (A) WT level current production in DHNA (20 μg/mL) containing media was supported by all conditions (*ΔeetA/B, ΔdmkA, ΔdmkB*) except *Δndh2* and *ΔpplA*. (B) *Δndh1* was not required for DHNA-dependent current production. (C) Iron oxide nanoparticle reduction was assayed after 24h anaerobic incubation in DHNA containing media with WT, *Δndh2, ΔpplA, ΔeetA/B, ΔdmkA, ΔdmkB. Δndh2* and *ΔpplA* have the largest decrease in reduced iron, followed by *ΔeetA/B* and then *ΔdmkA, ΔdmkB*. (D) *Δndh1* was not required for DHNA-dependent iron reduction. All Error shown is SD. Statistical significance calculated by t-test. *= p<0.05; ***= p<0.001

Having established the role of FLEET genes in DHNA-dependent anode reduction, we next sought to address the involvement of these genes in DHNA-dependent iron reduction. To understand which FLEET genes are required to reduce iron, we tested each mutant’s ability to reduce iron oxide nanoparticles under anaerobic conditions. Consistent with previous work, Δ*ndh2* and Δ*pplA* both showed a significant decrease in iron reduction compared to WT cells (Figure 3C) while Δ*dmkA* and Δ*dmkB* exhibited a minor, but significant decrease in reduced iron, while *ΔeetA/B* cells were found to have a slightly larger decrease in reduced iron compared to WT (Figure 3C). As seen in anode reduction, Δ*ndh1* cells were found to have no significant change in iron reduction compared to WT in the presence of DHNA (Figure 3D). Together, these data indicate that extracellular electron transfer begins at Ndh2 via the oxidation of cytosolic NADH. Electrons can then be directly transferred to extracellular DHNA or be transferred to PplA via EetA; ultimately reducing ferric iron or an anode (Figure 6; Table 2).

**Table 2:** Genetic requirements of EET activity.

|  | DHNA-dependent EET |  | Flavin-dependent EET |  |
| --- | --- | --- | --- | --- |
|  | Iron reduction | Anode reduction | Iron reduction | Anode reduction |
| *ndh2* | + | + | + | + |
| *eetA* or *eetB* | + | - | + | + |
| *pplA* | + | + | - | + |
| *ndh1* | - | - | - | - |
| *dmkA* | - | - | - | - |
| *dmkB* | - | - | - | - |

521 (+) required; (-) not required

### Riboflavin mediates electron transfer through an unknown, protein mediated mechanism

Having established the DHNA-dependent EET pathway in *L. plantarum*, we turned to interrogate the flavin-dependent EET pathway. The differential responses to DHNA and riboflavin we observed suggest that DHNA could be acting as an electron shuttle between the microbe and the electrode, while riboflavin likely requires a longer exposure to elicit its effect. To determine if riboflavin is acting as a mediator during flavin-dependent EET, we performed cyclic voltammetry on media containing freshly dissolved riboflavin, riboflavin that had been in the reactor for 24h, and riboflavin incubated with cells for 24 h. Freshly dissolved riboflavin has two prominent redox peaks at -493 mV and -416mV (Figure 4A). Interestingly, 24 hours on a poised electrode under microaerobic conditions resulted in the loss of the -400mV peak while the peak at -510 mV persisted (Figure 4A). Conversely, the presence of cells resulted in the loss of the peak at -500 mV and increased the amplitude of the redox peak at -399 mV (Figure 4A). This biotic shift suggests that cells may be converting riboflavin to FMN similar to results seen in previous studies.^19^

**Figure 4:**
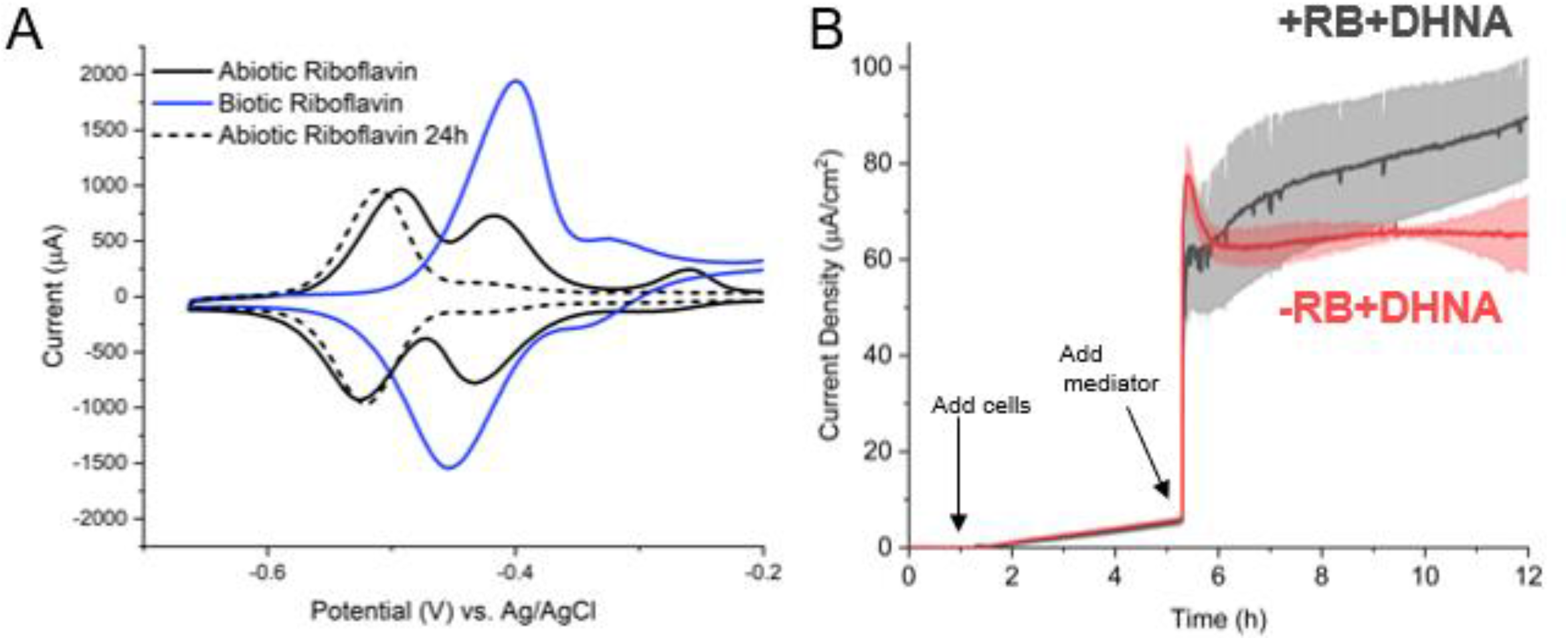
Riboflavin exhibits poor redox shuttle characteristics. (A) Cyclic voltammetry was performed prior to the addition of cells and 24h after the addition of cells. A shift right shift is observed for riboflavin in the presence of cells (B) *L. plantarum* was grown in microaerobic 3-chamber bioelectrochemical reactors for 3 hours on an electrode poised at 0.2V vs Ag/AgCl (3M KCl) before DHNA (red) or DHNA and riboflavin (gray) was injected. Current density measurements were taken every 36s for 12h. Error bars show SD.

Conversion of riboflavin to FMN by *L. plantarum* would require more time than immediate utilization of riboflavin. To interrogate the time-scale of response to the addition of riboflavin, we performed chronoamperometry following the addition of riboflavin+DHNA and DHNA alone. We found that the addition of riboflavin+DHNA did not result in an immediate increase in current compared to DHNA alone, but it did enable a greater maximum current density that was reached at ∼16h (Figure 4B). This finding strongly suggests that riboflavin does not act immediately to support anode reduction, but rather enhances EET by an unknown mechanism that requires time to incorporate, transport, or modify riboflavin.

We next sought to characterize the degree to which concentration can affect riboflavin’s ability to support EET using pulse-chase chronoamperometry. We found that the maximum current density increased with the riboflavin concentration, and there was a proportional decrease in the time to ½ the max current (Table 1). Overall, riboflavin supported current production was less robust compared to DHNA-dependent EET. Interestingly, the increase in riboflavin concentration did not result in a proportional increase in current density suggesting that cells may only utilize a particular concentration. In summary, it is likely that riboflavin could be supporting EET in part, as an electron shuttle, or as some form of cofactor.

To probe if riboflavin can serve as an electron shuttle, we performed another media swap using a graphite rod electrode, similar to the approach used with DHNA, to select for mediated electron transfer (MET). We found that riboflavin did support current production and that the current was dependent on the presence of riboflavin (SI Figure 4). These data suggest that riboflavin can serve as a shuttle under MET favorable conditions, but the primary mechanism by which riboflavin supports EET is likely by enhancing direct electron transfer rather than by acting as a mediator (Figure 6).

### Flavin-mediated EET requires Ndh2, PplA, and EetA

Based on our findings that riboflavin can support robust anode reduction in *L. plantarum*, we next sought to identify which FLEET genes were required to mediate flavin-dependent anode reduction. First, genetic knockouts were tested for their ability to reduce an anode. We observed no difference in current production between WT cells and *ΔdmkA* or *ΔdmkB* cells (Figure 5A). Alternatively, we observed a profound reduction in current production of *ΔpplA, Δndh2*, and *ΔeetA/B* compared to WT (Figure 5A). Because of potential side reactions with Ndh1, the non-FLEET NADH dehydrogenase, we tested *Δndh1* cells for their ability to produce current. Surprisingly, we observed the cells more quickly reach a higher max current density than the WT cells (Figure 5B). This increase in current density observed in *Δndh1* is likely because Ndh1 maintains redox balance and *Δndh1* cells must redox balance via Ndh2.Together, we show that flavin-dependent anode reduction precedes through Ndh2 to EetA and then to PplA in a riboflavin dependent manner. We propose that riboflavin can accept electrons from PplA via the FMN-ylated residues and ultimately to the electrode (Figure 6).

**Figure 5.**
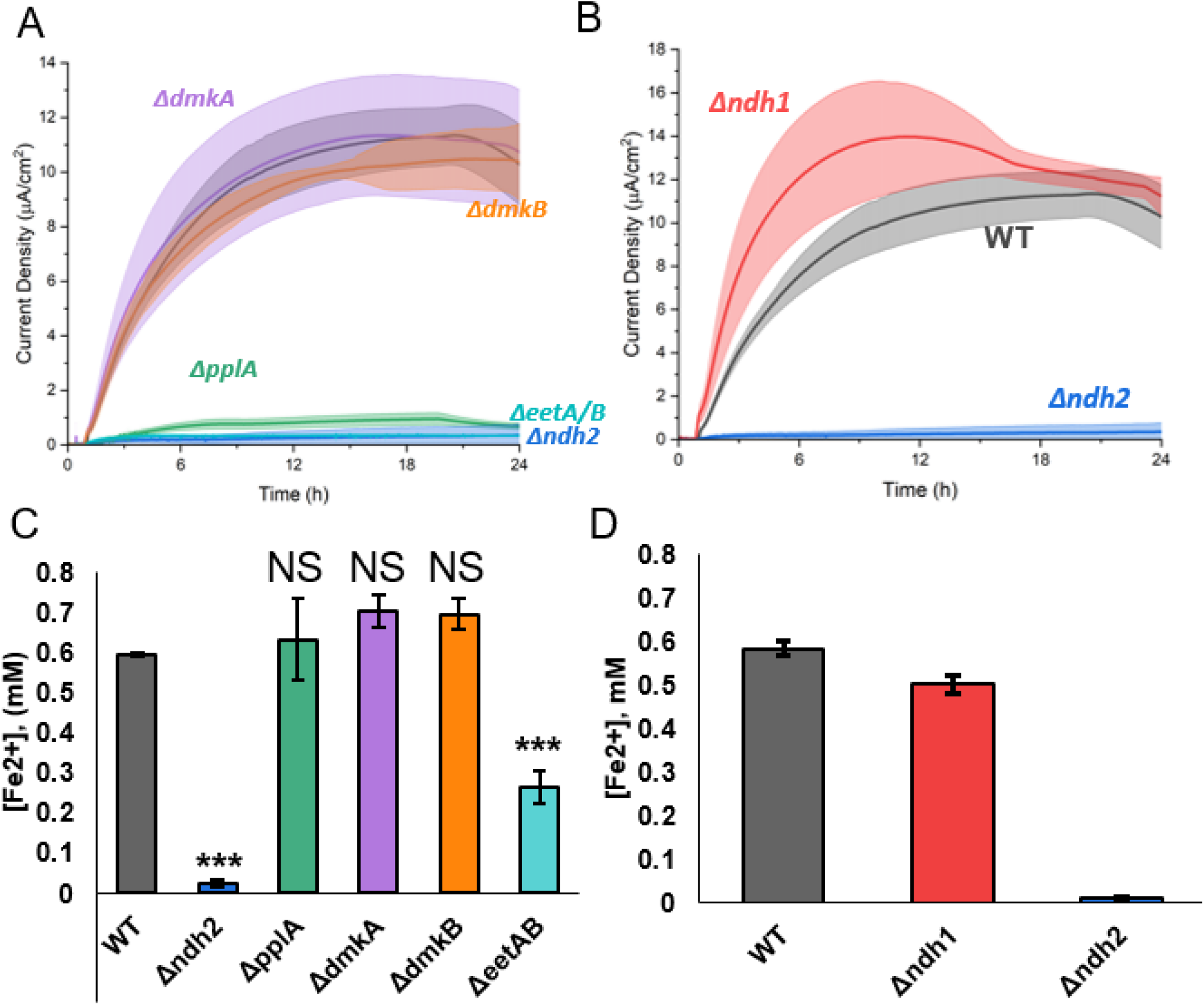
Flavin-dependent EET utilizes Ndh2, EetA, and PplA to reduce extracellular electron acceptors. Chronoamperometry was performed in 2-chambered 3 electrode bioreactors. (A) WT level current production in riboflavin (2 μg/mL) containing media was supported by *ΔdmkA* and *ΔdmkB*, while *Δndh2, ΔeetA/B*, and *ΔpplA* were unable to produce current. (B) *Δndh1* was not required for riboflavin-dependent current production and resulted in improved initial current production. (C) Iron oxide nanoparticle reduction was assayed after 24h anaerobic incubation in DHNA containing media with WT, *Δndh2, ΔpplA, ΔeetA/B, ΔdmkA, ΔdmkB. Δndh2. ΔpplA, ΔdmkA*, and *ΔdmkB* showed no change in reduced iron, while *ΔeetA/B* and *Δndh2* had significantly less reduced iron compared to WT. (D) *Δndh1* was not required for Flavin-dependent iron reduction.. All Error shown is SD. Statistical significance calculated by t-test. *= p<0.05; ***= p<0.001

**Figure 6:**
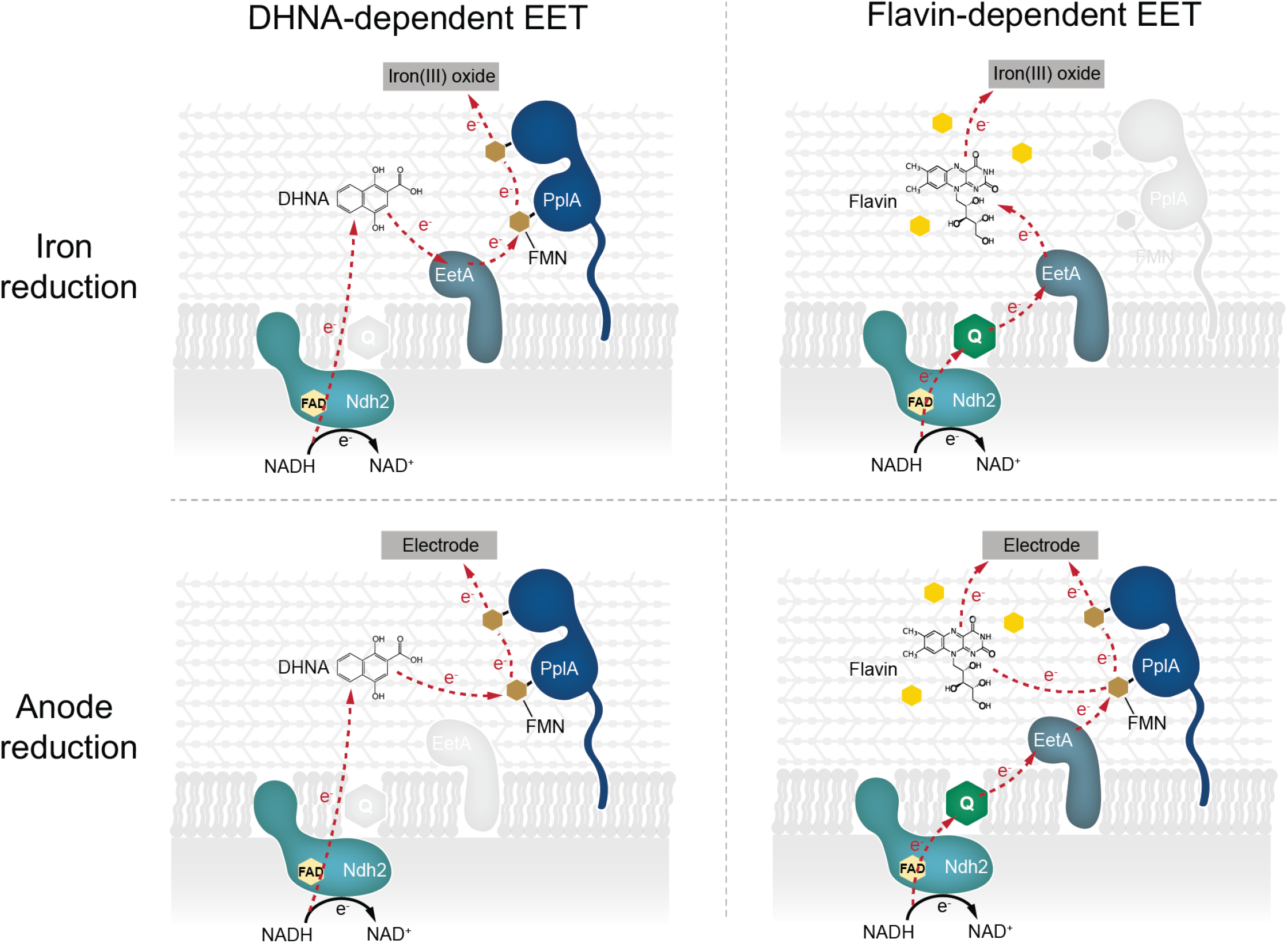
Schematic representation of DHNA- and Flavin-dependent EET routes. Taken together, this study found that EET can be supported by the presence of exogenous DHNA or Flavin species via two distinct routes. Moreover, the two routes appear to be optimized for different time scales and terminal electron acceptors.

With the characterization of the role of FLEET genes in flavin-dependent anode reduction, we next sought to address the involvement of these genes in flavin-dependent iron reduction. To assess the effect of genetic knockouts on electron transfer to extracellular iron, we measured iron reduction of FLEET knockout strains in the presence of riboflavin. Interestingly, we found that riboflavin could not support iron reduction in *Δndh2* or *ΔeetA/B* cells, but riboflavin could support iron reduction in *ΔpplA, ΔdmkA, ΔdmkB* and WT cells (Figure 5C). The lack of effect observed in the *ΔpplA* is surprising because it has such a strong phenotype under anode reduction conditions. We hypothesize that the PplA is not required for flavin-dependent iron reduction due to the presence of an unknown oxidoreductase or an unknown route of extracellular electron transfer to ferric oxides. Consistent with reactor conditions, *Δndh1* cells were found to have no significant change in iron reduction compared to WT in the presence of riboflavin (Figure 5D). Furthermore, we propose a novel route of flavin-based electron transfer that initiates from the oxidation of NADH by Ndh2 and proceeds through EetA to extracellular flavin to reduce insoluble iron oxide. Alternatively, in the case of an anode, electrons are transferred to PplA and ultimately to the anode (Figure 6; Table 2).

## Discussion

This study investigated the relative importance and mechanism of riboflavin and quinones for EET in *L. plantarum*. We found that both DHNA and riboflavin can independently support EET under minimal media conditions, but that riboflavin requires pregrowth with a quinone. Mechanistically, we observed that DHNA functions as a robust extracellular electron shuttle that likely receives its electrons from Ndh2 or PplA. Conversely, we found that riboflavin can also function as an electron shuttle - albeit less efficiently - and that it is likely being converted to FMN by cells to support EET to an anode. Moreover, we find that Ndh2, PplA, and EetA are required for anode reduction via riboflavin, but PplA is not required to reduce iron. Together, this study highlights novel routes of electron transfer that depend on both the redox mediator and the terminal electron acceptor (redox potential and accessibility) (Figure 6).

This work illuminates several aspects of flavin-dependent EET, but leaves some questions open. While one goal for this study was to minimize differences in cellular status before assaying iron and anode reduction, we still are left with a difference between flavin-dependent iron and anode reduction. This difference is relatively surprising as riboflavin facilitates reduction of iron and an electrode is kinetically favorable, as evidenced by a notable difference in midpoint potentials.^26^ We hypothesize this difference could be due to two variables, the nature of the terminal electron acceptor or the bioavailability of riboflavin. One such side reaction could be the result of the reactor setup we use for these experiments as it is maintained at a microaerobic state by N_2_ bubbling, but riboflavin has been shown to donate electrons to oxygen at the oxic-anoxic interface in other organisms.^27^ Finally, it is entirely possible that PplA is not required for flavin-dependent iron reduction due the functioning of a uncharacterized extracellular flavinated reductase powered by EET.

As a nomadic species that inhabits plant leaves, food/feed, soil and the gastrointestinal tract of various mammals, *L. plantarum* must adapt to diverse environmental niches.^28^ The identification of EET activity in LABs by this study and others poses the question, why would an iron-tolerant and great fermenter maintain an EET pathway? One explanation is that *L. plantarum* may utilize multiple routes as a means of increasing the likelihood of the microbe to perform EET and thus, enhance fitness.^9^ Additionally, *L. plantarum*, among other *Lactobacilli* species, exhibit niche-specific genetic alterations that have been linked to this organism’s ability to alter its metabolism and regulate the composition of the surrounding microbial community. ^29-31^ This study characterized the ability of *L. plantarum* to utilize biomolecules commonly released by microbes to support EET via overlapping, but unique mechanisms. We assert that multiple EET routes have been maintained by *L. plantarum* to support survival in diverse environments and which route is preferred is dependent on the availability of redox active molecules in the local environment and the cellular status (i.e. flavin-dependent EET prior exposure to a quinone to increase the electron flux). Moreover, this study begins to characterize the metabolic hierarchy utilized by *L. plantarum* to quickly adapt to local conditions.

Multiple studies have highlighted the probiotic benefits of *Lactiplantibacillus plantarum* such as, reducing host epithelial inflammation, altering host lipid metabolism, and exhibiting antihyperglycemic potential.^32–34^ Additionally, it is increasingly evident that microbes in the gut microbiome share and differentially utilize secreted factors like quinones and flavins to survive.^35^ We believe that the discovery of EET supported by flavins is of significant importance to understanding the dynamics of interspecies microbial energetics within the gut microbial community. EET activity can have a significant impact on microbial gut colonization as evidenced by a decreased CFU of EET mutants in studies of *L. monocytogenes* and *E. faecalis*.^6,36^ Moreover, since EET can facilitate increased fitness and colonization, this suggests that EET-active probiotics could have increased effectiveness. It appears that utilizing both DHNA- and flavin-dependent EET routes could enable *L. plantarum* to persist and thrive in complex environments and gain an energetic advantage over non-electrogenic microbes.

## Methods

### Microbial Cultivation

*Lactiplantibacillus plantarum* NCIMB8826 was obtained from Dr. Maria Marco. Bacterial strains were grown in MRS (HiMedia Laboratories) from glycerol stocks overnight at 37°C. Fresh overnight cultures were inoculated in mMRS(mannitol as carbon source) at an OD600 of ∼0.1 and grown for 16-18 h at 37°C. The composition of mMRS is as follows unless otherwise noted: 20 mg/mL D-mannitol (Sigma-Aldrich), 1% Tween 80 (Sigma-Aldrich), 10 mg/mL protease peptone (Gibco), 5 mg/mL yeast extract (Sigma-Aldrich), 11.48 mM dibasic potassium phosphate (Avantor Performance Materials), 61 mM sodium acetate trihydrate (Sigma-Aldrich), 8.83 mM tribasic ammonium citrate (Alfa Aesar), 0.83 mM anhydrous magnesium sulfate (Sigma-Aldrich), 0.3 mM manganese sulfate monohydrate (Sigma-Aldrich), 2 mM ferric ammonium citrate (Sigma-Aldrich), and 20 μg/mL 1,4-dihydroxy-2-naphthoic acid (DHNA, Sigma-Aldrich). Culture medium was supplemented with 10 μg/mL erythromycin (Sigma-Aldrich), or 10 μg/mL chloramphenicol (Sigma-Aldrich) when specified.

### Creation of gene deletions in L. plantarum NCIMB8826

The strains, plasmids, primers and DNA fragments used in this study are listed in Supplementary Table 1.

*L. plantarum* NCIMB8826 wild type, pplA, and ndh2 deletion mutant were kindly provided to us by Maria Marco.^9^ *L. plantarum* NCIMB8826 *dmkA, dmkB, ndh1* deletion mutants were constructed by CRISPR-Cas9 toolbox according to Huang et al.^25^ Briefly, the upstream and downstream homologous arms were amplified from genomic DNA of *L. plantarum* (see Supplementary Table 1 for primers), along with sgRNA fragments. Both fragments were cloned in ApaI-XbaI digested pHSP02 editing plasmid to create pSL08 (for *dmkA* ko), pSTS04 (for *dmkB* ko), pSL47(for *ndh1* ko) by Gibson assembly.^37^ For CRISPR editing, *L. plantarum* strain harboring helper plasmid pLH01 was induced with 100 ng/ml Sakain P peptide (GenScript, Piscataway, NJ) to express RecE/T and made electrocompetent. The editing plasmids were then delivered into *L. plantarum* NCIMB8826 by electroporation. The transformed cells were spread on MRS plates containing 10 μg/mL erythromycin and 10 μg/mL chloramphenicol to screen for the deletion mutants. Colony PCR was performed on single colonies to confirm the deletion of the target gene (see Supplementary Table 1 for primers). Following deletion confirmation, the plasmid was removed via serial growth in non-selective media.

### Assaying reduction of iron (III) oxide nanoparticles

To monitor their ability to reduce iron (III), 3 mL cultures of *L. plantarum* grown in mMRS were grown overnight. The cells from these cultures were pelleted by centrifugation at 4000x*g* for 10 min at 4°C and washed with PBS two times. The washed cells were resuspended in PBS to an OD600 of 2.0. These cell suspensions were combined with an equal volume of assay master mix to yield samples with a final volume of 0.5-1 mL. The assay master mix contained 40 mg/mL D-mannitol, 40 μg/mL DHNA, 4 mM ferric oxide (<50 nm nanoparticle, Sigma Aldrich), and 2X PBS (pH= 7.4).The samples were introduced into an anaerobic chamber maintained between 30 °C to 34 °C and were incubated for 24 h before iron reduction was measured. To prevent oxidation of iron upon oxygen exposure, samples were diluted 1:1 in 0.5 M HCl while in the anaerobic chamber. After removal from the anaerobic chamber, the samples were centrifuged at 4000 x g for 10 min at 4°C, and the cell-free supernatant was collected. Fifty μL of supernatant was added to 200 μL of 2 mM ferrozine in HEPES buffered to pH 7. After 5 min of incubation in the dark, the absorbance was measured at 562 nm. Iron concentration was calculated by comparison of the measured A_562nm_ to a standard curve of FeSO_4_ ranging from 0.4-0.025 mM.

### Bioelectrochemical measurements

All bioelectrochemical measurements were performed on VSP-300 potentiostat (Biologic). Unless otherwise noted, all measurements were performed using a carbon-felt (Alfa Aesar) platinum electrode (Alfa Aesar) poised at 0.2V vs Ag/AgCl in 3M KCl for chronoamperometry. Current measurements were collected every 36 s for the duration of the experiment. Cyclic voltammetry was performed from -0.7 to 0.7V at a scan rate of 5 mV/s. We used a two-chambered, 3 electrode set-up separated by a cationic membrane. The working chamber contained filter sterilized PBS (pH7.4) and 20 mg/mL Mannitol. The media was supplemented with 20 μg/mL DHNA, 2μg/mL riboflavin, or both when specified.

### Media Swap Experiment

Cells were grown and washed as noted previously before being injected into bioelectrochemical reactors containing PBS, 20 mg/mL mannitol. A graphite rod electrode was poised at 0.2V vs Ag/AgCl and chronoamperometric measurements were collected every 36s. After ∼3h, current production leveled out and 2 µg/mL riboflavin or 20 µg/mL DHNA was added. After the peak current was achieved and maintained (∼14h), cells were collected, washed and kept on ice until reactors with fresh media without riboflavin were purged. Cells were injected into the new reactors and chronoamperometry was performed. After ∼18h, 2 μg/mL riboflavin or 20 µg/mL DHNA was added back to the reactors.

## Literature Cited

(1) Chen, L.; Cao, C.; Wang, S.; Varcoe, J. R.; Slade, R. C. T.; Avignone-Rossa, C.; Zhao, F. Electron Communication of Bacillus Subtilis in Harsh Environments. iScience 2019, 12, 260–269. https://doi.org/10.1016/j.isci.2019.01.020.

(2) Shi, L.; Dong, H.; Reguera, G.; Beyenal, H.; Lu, A.; Liu, J.; Yu, H.-Q.; Fredrickson, J. K. Extracellular Electron Transfer Mechanisms between Microorganisms and Minerals. Nat. Rev. Microbiol. 2016, 14 (10), 651–662. https://doi.org/10.1038/nrmicro.2016.93.

(3) Paquete, C. M. Electroactivity across the Cell Wall of Gram-Positive Bacteria. Comput. Struct. Biotechnol. J. 2020, 18, 3796–3802. https://doi.org/10.1016/j.csbj.2020.11.021.

(4) Okamoto, A.; Nakamura, R.; Hashimoto, K. In-Vivo Identification of Direct Electron Transfer from Shewanella Oneidensis MR-1 to Electrodes via Outer-Membrane OmcA–MtrCAB Protein Complexes. Electrochimica Acta 2011, 56 (16), 5526–5531. https://doi.org/10.1016/j.electacta.2011.03.076.

(5) Light, S. H.; Su, L.; Rivera-Lugo, R.; Cornejo, J. A.; Louie, A.; Iavarone, A. T.; Ajo-Franklin, C. M.; Portnoy, D. A. A Flavin-Based Extracellular Electron Transfer Mechanism in Diverse Gram-Positive Bacteria. Nature 2018, 562 (7725), 140–144. https://doi.org/10.1038/s41586-018-0498-z.

(6) Light, S. H.; Méheust, R.; Ferrell, J. L.; Cho, J.; Deng, D.; Agostoni, M.; Iavarone, A. T.; Banfield, J. F.; D’Orazio, S. E. F.; Portnoy, D. A. Extracellular Electron Transfer Powers Flavinylated Extracellular Reductases in Gram-Positive Bacteria. Proc. Natl. Acad. Sci. 2019, 116 (52), 26892–26899. https://doi.org/10.1073/pnas.1915678116.

(7) Keogh, D.; Lam, L. N.; Doyle, L. E.; Matysik, A.; Pavagadhi, S.; Umashankar, S.; Williams, R. B. H.; Marsili, E.; Kline, K. A. Extracellular Electron Transfer Powers Enterococcus Faecalis Biofilm Metabolism. 2018, 9 (2), 17.

(8) Pankratova, G.; Leech, D.; Gorton, L.; Hederstedt, L. Extracellular Electron Transfer by the Gram-Positive Bacterium Enterococcus Faecalis. Biochemistry 2018, 57 (30), 4597–4603. https://doi.org/10.1021/acs.biochem.8b00600.

(9) Tejedor-Sanz, S.; Stevens, E. T.; Li, S.; Finnegan, P.; Nelson, J.; Knoesen, A.; Light, S. H.; Ajo-Franklin, C. M.; Marco, M. L. Extracellular Electron Transfer Increases Fermentation in Lactic Acid Bacteria via a Hybrid Metabolism. eLife 2022, 11, e70684. https://doi.org/10.7554/eLife.70684.

(10) Francis, I.; Holsters, M.; Vereecke, D. The Gram-Positive Side of Plant–Microbe Interactions. Environ. Microbiol. 2010, 12 (1), 1–12. https://doi.org/10.1111/j.1462-2920.2009.01989.x.

(11) Barbosa, A. A. T.; Mantovani, H. C.; Jain, S. Bacteriocins from Lactic Acid Bacteria and Their Potential in the Preservation of Fruit Products. Crit. Rev. Biotechnol. 2017, 37 (7), 852–864. https://doi.org/10.1080/07388551.2016.1262323.

(12) Thakur, K.; Tomar, S. K.; De, S. Lactic Acid Bacteria as a Cell Factory for Riboflavin Production. Microb. Biotechnol. 2015, 9 (4), 441–451. https://doi.org/10.1111/1751-7915.12335.

(13) Mathur, H.; Beresford, T. P.; Cotter, P. D. Health Benefits of Lactic Acid Bacteria (LAB) Fermentates. Nutrients 2020, 12 (6), 1679. https://doi.org/10.3390/nu12061679.

(14) Li, X.; Liu, L.; Liu, T.; Yuan, T.; Zhang, W.; Li, F.; Zhou, S.; Li, Y. Electron Transfer Capacity Dependence of Quinone-Mediated Fe(III) Reduction and Current Generation by Klebsiella Pneumoniae L17. Chemosphere 2013, 92 (2), 218–224. https://doi.org/10.1016/j.chemosphere.2013.01.098.

(15) Lin, X.; Yang, F.; You, L.; Wang, H.; Zhao, F. Liposoluble Quinone Promotes the Reduction of Hydrophobic Mineral and Extracellular Electron Transfer of Shewanella Oneidensis MR-1. The Innovation 2021, 2 (2), 100104. https://doi.org/10.1016/j.xinn.2021.100104.

(16) Fuller, S. J.; McMillan, D. G. G.; Renz, M. B.; Schmidt, M.; Burke, I. T.; Stewart, D. I. Extracellular Electron Transport-Mediated Fe(III) Reduction by a Community of Alkaliphilic Bacteria That Use Flavins as Electron Shuttles. Appl. Environ. Microbiol. 2014, 80 (1), 128–137. https://doi.org/10.1128/AEM.02282-13.

(17) Marsili, E.; Baron, D. B.; Shikhare, I. D.; Coursolle, D.; Gralnick, J. A.; Bond, D. R. Shewanella Secretes Flavins That Mediate Extracellular Electron Transfer. Proc. Natl. Acad. Sci. 2008, 105 (10), 3968–3973. https://doi.org/10.1073/pnas.0710525105.

(18) Kotloski, N. J.; Gralnick, J. A. Flavin Electron Shuttles Dominate Extracellular Electron Transfer by Shewanella Oneidensis. mBio 4 (1), e00553–12. https://doi.org/10.1128/mBio.00553-12.

(19) Okamoto, A.; Hashimoto, K.; Nealson, K. H.; Nakamura, R. Rate Enhancement of Bacterial Extracellular Electron Transport Involves Bound Flavin Semiquinones. Proc. Natl. Acad. Sci. 2013, 110 (19), 7856–7861. https://doi.org/10.1073/pnas.1220823110.

(20) Okamoto, A.; Saito, K.; Inoue, K.; Nealson, K. H.; Hashimoto, K.; Nakamura, R. Uptake of Self-Secreted Flavins as Bound Cofactors for Extracellular Electron Transfer In Geobacter Species. Energy Env. Sci 2014, 7 (4), 1357–1361. https://doi.org/10.1039/C3EE43674H.

(21) Gurumurthy, D. M.; Bharagava, R. N.; Kumar, A.; Singh, B.; Ashfaq, M.; Saratale, G. D.; Mulla, S. I. EPS Bound Flavins Driven Mediated Electron Transfer in Thermophilic Geobacillus Sp. Microbiol. Res. 2019, 229, 126324. https://doi.org/10.1016/j.micres.2019.126324.

(22) Hederstedt, L.; Gorton, L.; Pankratova, G. Two Routes for Extracellular Electron Transfer in Enterococcus Faecalis. J. Bacteriol. 2020. https://doi.org/10.1128/JB.00725-19.

(23) Mevers, E.; Su, L.; Pishchany, G.; Baruch, M.; Cornejo, J.; Hobert, E.; Dimise, E.; Ajo-Franklin, C. M.; Clardy, J. An Elusive Electron Shuttle from a Facultative Anaerobe. eLife 2019, 8, e48054. https://doi.org/10.7554/eLife.48054.

(24) Matsui, Y.; Hamamoto, K.; Kitazumi, Y.; Shirai, O.; Kano, K. Diffusion-Controlled Mediated Electron Transfer-Type Bioelectrocatalysis Using Microband Electrodes as Ultimate Amperometric Glucose Sensors. Anal. Sci. 2017, 33 (7), 845–851. https://doi.org/10.2116/analsci.33.845.

(25) Huang, H.; Song, X.; Yang, S. Development of a RecE/T-Assisted CRISPR-Cas9 Toolbox for Lactobacillus. Biotechnol. J. 2019, 14 (7), e1800690. https://doi.org/10.1002/biot.201800690.

(26) Shi, Z.; Zachara, J. M.; Shi, L.; Wang, Z.; Moore, D. A.; Kennedy, D. W.; Fredrickson, J. K. Redox Reactions of Reduced Flavin Mononucleotide (FMN), Riboflavin (RBF), and Anthraquinone-2,6-Disulfonate (AQDS) with Ferrihydrite and Lepidocrocite. Environ. Sci. Technol. 2012, 46 (21), 11644–11652. https://doi.org/10.1021/es301544b.

(27) Khan, M. T.; Duncan, S. H.; Stams, A. J. M.; van Dijl, J. M.; Flint, H. J.; Harmsen, H. J. M. The Gut Anaerobe Faecalibacterium Prausnitzii Uses an Extracellular Electron Shuttle to Grow at Oxic–Anoxic Interphases. ISME J. 2012, 6 (8), 1578–1585. https://doi.org/10.1038/ismej.2012.5.

(28) Liu, W., Pang, H., Zhang, H., Cai, Y. (2014). Biodiversity of Lactic Acid Bacteria. In: Zhang, H., Cai, Y. (eds) Lactic Acid Bacteria. Springer, Dordrecht. https://doi.org/10.1007/978-94-017-8841-0_2

(29) Pan, M.; Hidalgo-Cantabrana, C.; Barrangou, R. Host and Body Site-Specific Adaptation of Lactobacillus Crispatus Genomes. NAR Genomics and Bioinformatics 2020, 2 (1), nqaa001. https://doi.org/10.1093/nargab/lqaa001.

(30) Pan, Q.; Cen, S.; Yu, L.; Tian, F.; Zhao, J.; Zhang, H.; Chen, W.; Zhai, Q. Niche-Specific Adaptive Evolution of Lactobacillus Plantarum Strains Isolated From Human Feces and Paocai. Frontiers in Cellular and Infection Microbiology 2021, 10.

(31) Cen, S.; Yin, R.; Mao, B.; Zhao, J.; Zhang, H.; Zhai, Q.; Chen, W. Comparative Genomics Shows Niche-Specific Variations of Lactobacillus Plantarum Strains Isolated from Human, Drosophila Melanogaster, Vegetable and Dairy Sources. Food Bioscience 2020, 35, 100581. https://doi.org/10.1016/j.fbio.2020.100581.

(32) Youn, H. S.; Kim, J.-H.; Lee, J. S.; Yoon, Y. Y.; Choi, S. J.; Lee, J. Y.; Kim, W.; Hwang, K. W. Lactobacillus Plantarum Reduces Low-Grade Inflammation and Glucose Levels in a Mouse Model of Chronic Stress and Diabetes. Infect. Immun. 89 (8), e00615–20. https://doi.org/10.1128/IAI.00615-20.

(33) Li, H.; Liu, F.; Lu, J.; Shi, J.; Guan, J.; Yan, F.; Li, B.; Huo, G. Probiotic Mixture of Lactobacillus Plantarum Strains Improves Lipid Metabolism and Gut Microbiota Structure in High Fat Diet-Fed Mice. Front. Microbiol. 2020, 11.

(34) Zhong, H.; Abdullah, Zhang Y.; Zhao, M.; Zhang, J.; Zhang, H.; Xi, Y.; Cai, H.; Feng, F. Screening of Novel Potential Antidiabetic Lactobacillus Plantarum Strains Based on in Vitro and in Vivo Investigations. LWT 2021, 139, 110526. https://doi.org/10.1016/j.lwt.2020.110526.

(35) Daisley, B. A.; Koenig, D.; Engelbrecht, K.; Doney, L.; Hards, K.; Al, K. F.; Reid, G.; Burton, J. P. Emerging Connections between Gut Microbiome Bioenergetics and Chronic Metabolic Diseases. Cell Rep. 2021, 37 (10), 110087. https://doi.org/10.1016/j.celrep.2021.110087.

(36) Lam, L. N.; Wong, J. J.; Matysik, A.; Paxman, J. J.; Chong, K. K. L.; Low, P. M.; Chua, Z. S.; Heras, B.; Marsili, E.; Kline, K. A. Sortase-Assembled Pili Promote Extracellular Electron Transfer and Iron Acquisition in Enterococcus Faecalis Biofilm. bioRxiv April 7, 2019, p 601666. https://doi.org/10.1101/601666.

(37) Gibson, D. G.; Young, L.; Chuang, R.-Y.; Venter, J. C.; Hutchison, C. A.; Smith, H. O. Enzymatic Assembly of DNA Molecules up to Several Hundred Kilobases. Nat. Methods 2009, 6 (5), 343–345. https://doi.org/10.1038/nmeth.1318.

